# Transient definitive host presence is sufficient to sustain avian schistosome populations

**DOI:** 10.1101/2021.10.05.463157

**Authors:** Sydney P Rudko, Brooke A McPhail, Ronald L Reimink, Kelsey Froelich, Alyssa Turnbull, Patrick C Hanington

## Abstract

To control swimmer’s itch in northern Michigan inland lakes, one species of bird, the common merganser (*Mergus merganser*), has been relocated from several lakes since 2015. Relocation efforts are driven by a desire to reduce the prevalence of the swimmer’s itch-causing parasite *Trichobilharzia stagnicolae*. The intention of this state-sponsored control effort was to interrupt the life cycle of *T. stagnicolae* and reduce parasite egg contribution into the environment from summer resident mergansers such that infections of the intermediate snail host *Stagnicola emarginata* declined. Reduced snail infection prevalence was expected to greatly reduce abundance of the swimmer’s itch-causing cercarial stage of the parasite in water. With no official program in place to assess the success of this relocation effort, we sought to study the effectiveness and impact of the removal of a single definitive host from a location with high definitive host and parasite diversity. This was assessed through a comprehensive, lake-wide monitoring study measuring longitudinal changes in the abundance of three species of avian schistosome cercariae in four inland Michigan lakes. Environmental measurements were also taken at these lakes to understand how they can affect swimmer’s itch incidence. Results from this study demonstrate that the diversity of avian schistosomes at the study lakes would likely make targeting of a single species of swimmer’s itch-causing parasite meaningless from a swimmer’s itch control perspective. Our data also suggest that removal of the common merganser is not an effective control strategy for the *T. stagnicolae* parasite, likely due to parasite contributions of migratory birds in the fall and spring. This suggests that only minimal contact time between the definitive host and the lake ecosystem is required to contribute sufficient parasite numbers to maintain a thriving population of parasite species with a high host-specificity.

## Introduction

Schistosomes are parasitic flatworms that rely on two hosts to complete their life cycles. A gastropod intermediate host supports asexual reproduction of the parasite, and numerous species of vertebrate can serve as definitive hosts, where adult worms reside, and sexual reproduction occurs. Humans serve as definitive hosts for specific species of schistosomes that cause a condition known as schistosomiasis, which affects over 229 million people throughout the world [1]. A closely related group of schistosomes commonly use waterfowl as their definitive host. The larval stage (cercariae) of these avian schistosomes can accidentally penetrate human skin where they die and cause the condition cercarial dermatitis, also commonly known as swimmer’s itch.

For a schistosome to thrive, both hosts must be present in an ecosystem to facilitate life cycle completion. It is often assumed that definitive host removal (whether physically or via chemotherapy to eliminate parasite infections) will break the life cycle sufficiently to reduce parasite abundance in the environment [2,3]. For human trematodes, such as *Schistosoma mansoni*, or *S. haematobium*, this assumption has been partially tested with control programs that focus on treatment of the human host using anthelmintic drugs (Praziquantel) in the hope that this treatment would sufficiently limit transmission to the snail host, thereby reducing the likelihood of human reinfection [4]. Work with removing avian hosts at the ecosystem level in California showed mixed results, with some parasites increasing, some decreasing, and others unaffected when the definitive bird hosts were removed [5]. In Michigan, USA, a program that supported the complete relocation of summer resident common mergansers (*Mergus merganser*) from inland lakes [6] assumed that removal of the definitive host will interrupt the life cycle of the avian schistosome *Trichobilharzia stagnicolae* and reduce the occurrence of swimmer’s itch.

Common mergansers are currently the only described definitive host for the avian schistosome *Trichobilharzia stagnicolae* [7] and show the highest infection rate of all waterfowl surveyed in Michigan inland lakes [8,9]. Due to this high infection prevalence, common mergansers became the focus of much of the swimmer’s itch research in the area. Despite the presence of numerous other waterfowl species known to harbor avian schistosomes, the common merganser was targeted and is now often harassed, illegally shot, and generally loathed by many riparians [10]. The Michigan Department of Natural Resources (MDNR) approved a 3-year pilot research study on Higgins Lake (Roscommon Co., MI) in 2014 and another pilot study on 5 lakes in northern Michigan in 2017 to trap and relocate 400 common mergansers to predetermined relocation sites on Lake Michigan or Lake Huron [11]. It was believed that removing the summer resident birds from the lake would interrupt the life cycle of *T. stagnicolae* and decrease incidence of swimmer’s itch on these lakes [9–11]. These pilot studies gave birth to a nuisance control permit developed by the MDNR.

Swimmer’s itch incidence can also be affected by environmental variables that influence the concentration and location of the cercariae in the water. These variables include water temperature, wind speed, and water movement [2,3,12–14]. Temperature has been shown to affect cercarial emergence from the snail intermediate hosts, which results in more cercariae being released as the water warms [3]. Wind speeds and water movement may coincide, as they can concentrate the cercariae and move them to another part of the water body or accumulate the cercariae close to shore where recreators may be present [2,3,14]. These variables could also result in a higher chance of encountering swimmer’s itch-causing parasites at some areas around the lake than others. Understanding these patterns may help to reduce cases of swimmer’s itch as recreators become educated about environmental conditions that increase their chance of developing dermatitis.

Using qPCR-based cercariometry, we have been tracking the relative abundance of the three most common species of swimmer’s itch-causing avian schistosomes in inland lakes of northern Michigan since 2018. This species-specific analysis was complemented by qPCR quantification of total avian schistosome abundance measurements [15–18]. Relative abundance of *T. stagnicolae* on lakes where common merganser relocation had been ongoing indicated that this primary target of the relocation effort was still present, and even the dominant avian schistosome species in some cases [18]. Because there is no current evidence *T. stagnicolae* utilizes other species of waterfowl as a definitive host [7], we hypothesized common mergansers must still be contributing *T. stagnicolae* to the relocation lakes, likely originating from non-resident spring and fall migrating birds which are still utilizing inland Michigan lakes as they migrate.

To better understand how avian schistosome populations fluctuate over a season on lakes where common mergansers are actively removed and on lakes left in their natural state, we undertook a longitudinal study. This study was designed to observe and understand changes in abundance of three avian schistosome species using qPCR cercariometry. For two years, weekly water samples along with environmental measurements were collected between mid-May and late October at thirty independent sites on four lakes in 2019 and 2020 (Glen Lake, South Lake Leelanau and North Lake Leelanau in Leelanau County, and Walloon Lake in Charlevoix County) in northern Michigan [18]. This was a massive undertaking, made possible through the help of community partners. We utilized qPCR testing to measure the relative abundance of *T. stagnicolae, T. physellae*, and a recently discovered avian schistosome currently known as Avian Schistosome C [19]. Key findings of this study suggest that *T. stagnicolae* abundance is not substantially impacted by relocation of the common merganser definitive host. The two study lakes at which merganser relocation efforts had been implemented were both dominated by the parasite these efforts were attempting to impact. While the relative abundance of each avian schistosome species we monitored varied between study lakes, no lake was exclusively inhabited by only a single avian schistosome species.

Broadly, the results of this study have implications beyond swimmer’s itch control efforts. Efforts to reduce transmission of human schistosomiasis are predicated on a similar assumption that schistosome life cycles can be interrupted at the definitive host stage via mass drug administration efforts. Here, we show that transient definitive host contribution of eggs to an ecosystem are sufficient to sustain a substantial, lake-wide population of schistosomes. Results of this investigation may be of use for informing strategies to better implement human schistosome elimination efforts, which should be striving to prevent any sources of environmental egg contribution if life cycle interruption is a goal.

## Results

### Avian schistosome populations differed between each lake

Evaluating the study lakes based on the total avian schistosome population revealed clear distinctions between Glen Lake and Walloon Lake. Most notable, was that in both 2019 and 2020, avian schistosomes were detected in abundance at Walloon Lake three weeks earlier than at Glen Lake. This observation was obvious at most of the Walloon Lake sites. Additionally, the density of the avian schistosome population was found to be substantially higher in May and June at Walloon Lake when compared to Glen Lake. Such distinctions were not as clear when comparing North and South Lake Leelanau to each other, or to Glen or Walloon Lake (Figure 1; Figure S1).

**Figure 1:**
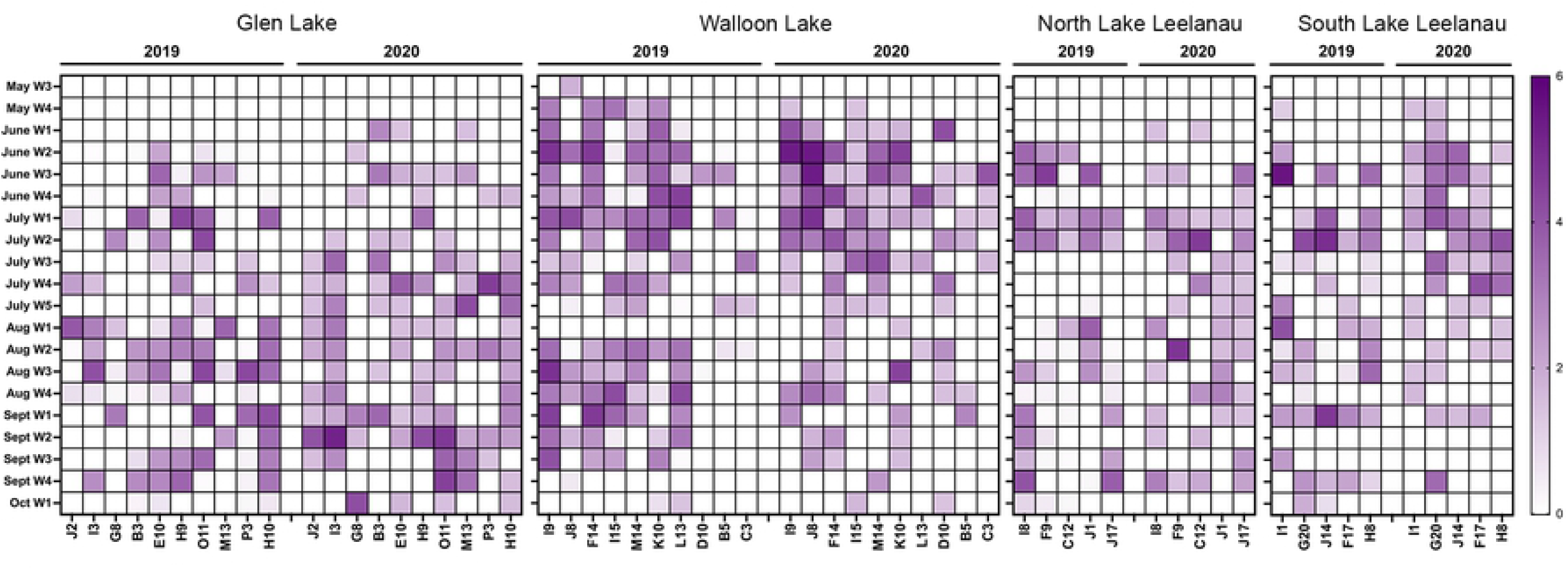
Log_10_-transformed estimates of total schistosome cercariae abundance (cercariae/25L lake water) organized by lake, sample site and date of sample collection. Relative intensity of the purple colour in each box reflects varying cercariae abundance. Absence of colour indicated no cercariae detected in the sample. Sample collection dates are identified by which week of the month they occurred.

At Glen Lake, 70 sites were positive throughout 2019 for avian schistosomes compared to 90 positive sites in 2020. Only 10% (7) and 14% (13) of the positive sites at Glen Lake were collected before the end of June in 2019 and 2020 respectively (Figure 1). This is contrasted by Walloon Lake, at which 33% (96) of sites and 90 (36%) of sites had detectable avian schistosomes prior to the end of June (Figure 1). A comparison of the mean estimates of cercariae number per 25 litres of water from all positive sites sampled at Glen (77.16 +/- SD 465.77) and Walloon (2975.07 +/- SD 9347.1) Lakes prior to the end of June indicated that there was a significant difference (p=0.02) despite large standard deviations. The same was true for the comparison in 2020, where mean estimates of avian cercariae number at Glen Lake (39.7 +/- SD 176.48) and Walloon Lake (13518.97 +/- SD 47491.66) yielded a statistically significant difference (p=0.04) as well (Figure 1).

The number of avian schistosome-positive sites prior to July 1 at North and South Lake Leelanau were similar in 2019 with 7 (23%) and 5 (17%) of sites. In 2020, North Lake Leelanau remained consistent with 5 (17%) positive sites before July 1, however, South Lake Leelanau increased the number of positive sites to 14 (47%).

### *T. stagnicolae* persists even at lakes with 3-year long control programs

Underpinning the higher-level distinctions between the four study lakes is the fact that the avian schistosome populations are comprised of very different species of avian schistosome. *T. stagnicolae* consistently dominated the avian schistosome population at Glen Lake and North Lake Leelanau in 2019 and 2020 despite sustained control programs for three years prior to 2019, which targeted the definitive merganser host (Figure 2, panels A and C). At Glen Lake *T. stagnicolae* was first detected in the water on June 11, 2019, and its presence was detected consistently until October 01, 2019. The frequency of detection was highly variable between sites. At some sites (notably E10, H10, H9, I3, O11, and P3) *T. stagnicolae* was detected consistently; however, at other sites, the parasite was rarely detected (i.e.: G8, M13) (Figure 2).

**Figure 2:**
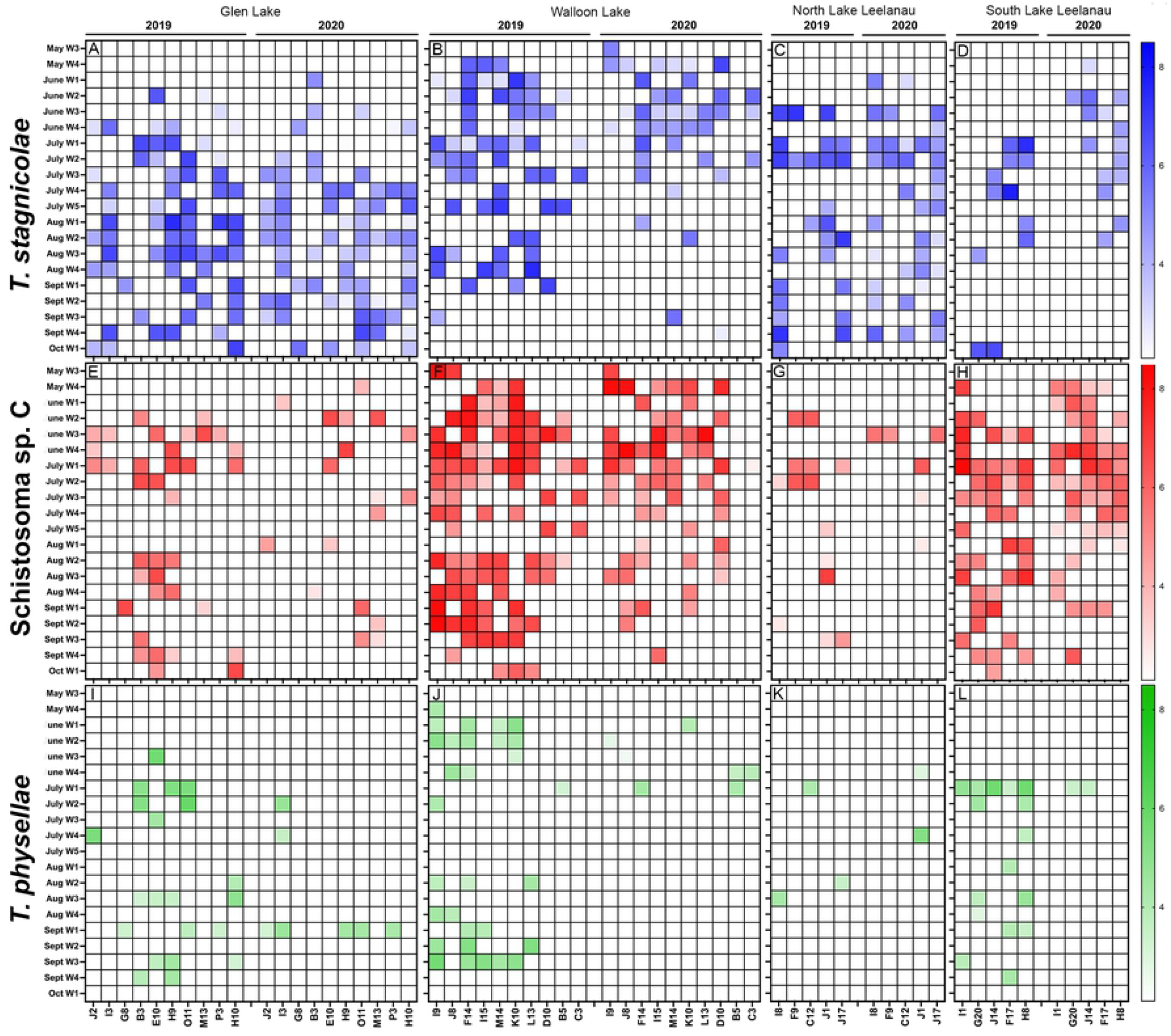
Log_10_-transformed estimates of target avian schistosome species CO1 gene copies organized by lake, sample site and date of sample collection. Relative intensity of the colour in each box reflects varying CO1 gene abundance. Absence of colour indicates that no cercariae were detected in the sample using the total schistosome qPCR test, and thus, these samples were not assessed using the species-specific tests. Sample collection dates are identified by which week of the month they occurred.

Lake-wide, *T. stagnicolae* signal rose sharply starting in July 2019 and 2020 at Glen Lake. However, at Walloon Lake and North Lake Leelanau, which both sustained consistent populations of *T. stagnicolae* through 2019 and 2020, the *T. stagnicolae* signal is detected earlier (late May-early June at Walloon Lake and early June at North Lake Leelanau). *T. stagnicolae* DNA abundance maintained consistently high concentrations continuing through August 2019 and September 2019 at Glen Lake and North Lake Leelanau until sharply dropping (Figure 2; Figure 3; Figure S1). The *T. stagnicolae* population at Walloon Lake tapers off earlier, with only a single positive site observed for both 2019 and 2020 after the end of August (Figure 2B).

**Figure 3:**
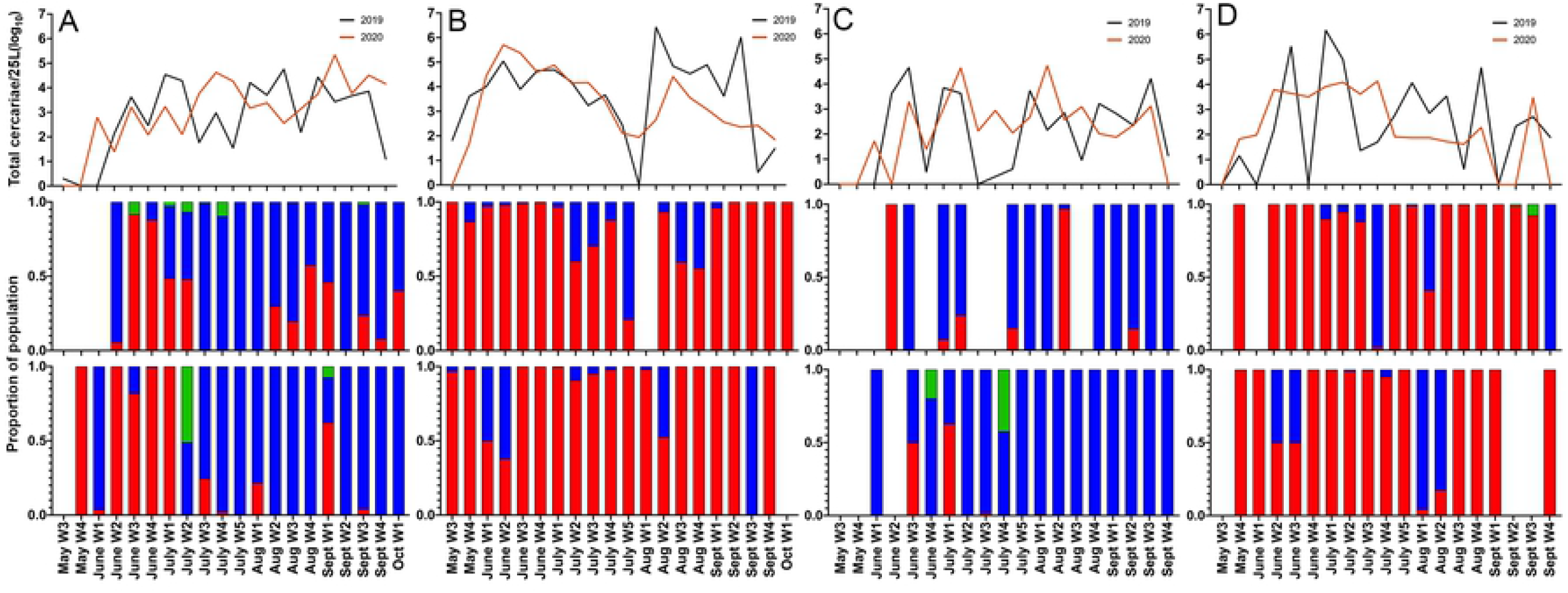
Accumulated Log_10_ transformed cercariae/25L for 2019 and 2020 shown for each study lake (A: Glen Lake; B: Walloon Lake; C: North Lake Leelanau; D: South Lake Leelanau) with the proportional contribution of each assessed avian schistosome species (red = Avian schistosome sp. C, blue = *T. stagnicolae*, green = *T. physellae*) for 2019 (top bar chart) and 2020 (lower bar chart).

*T. stagnicolae* was the dominant avian schistosome species at North Lake Leelanau in 2019 and 2020 (this lake implemented a merganser control program from 2017 ending in 2019). Similar to Glen Lake, *T. stagnicolae* signal increased sharply in late June, early July, and looking at the lake-wide accumulated data from all sites, stayed at between 6 and 7 log_10_ abundance until early October in both 2019 and 2020 (Figure 3; Figure S1). At the site-specific level, again similar to Glen Lake, some sites at North Lake Leelanau had a higher *T. stagnicolae* signal than others. Sites I8, J1 and J17 in 2019 all consistently have *T. stagnicolae* signal present, whereas sites C12 and F9 have little to no detectable *T. stagnicolae* DNA (Figure 2C).

South Lake Leelanau has never had a control program because no merganser broods have been observed at this lake since surveys began in 2016. For that reason, South Lake Leelanau served as a negative control for the presence of resident merganser broods. In 2019, South Lake Leelanau had high concentrations of *T. stagnicolae* DNA despite the absence of the summer resident definitive host, and lake wide data suggests that the abundance of this parasite was comparable to North Lake Leelanau (between 4.5-7.5 log_10_ gene copies (GC)/sample) (Figure 3). The site-specific data indicates a patchy distribution of *T. stagnicolae*-positive sites (Figure 2). Walloon lake served as a positive control lake since it has never had a merganser control program and has numerous yearly common merganser broods present. *T. stagnicolae* also had a high abundance at this lake in 2019 that rose sharply in early June and tapered off towards the end of September (Figure 2; Figure 3) and had a similarly “patchy” distribution in *T. stagnicolae* density, with some sites supporting more parasite than others.

In 2020, Glen Lake and North Lake Leelanau ceased relocating merganser broods to control swimmer’s itch. Detection and abundance of *T. stagnicolae* DNA at Glen Lake was similar between 2019 and 2020 (Figure 2; Figure 3). North Lake Leelanau also had an increase in the number of days cercariae are found in the water, and site J17, another site that in 2019 was relatively cercariae free now had detectable cercariae present in the water on nearly every sampling day (Figure 3). *T. stagnicolae* profiles at South Lake Leelanau and Walloon Lake appear relatively unchanged from 2019 to 2020; however, the presence of *T. stagnicolae* at both lakes is shifted to earlier in the spring/summer period, with few instances of *T. stagnicolae* being encountered in the water after July 21, 2020 at both lakes.

### Novel species Avian Schistosome C is an important contributor to avian schistosome populations in Northern Michigan

The newly discovered and yet to be named Avian schistosome C was the most abundant parasite at both Walloon and South Lake Leelanau in 2019 and 2020. It was less abundant than *T. stagnicolae* at Glen Lake and North Lake Leelanau in both years (Figure 3; Figure S1). South Lake Leelanau was the only lake that displayed a significant difference in Avian schistosome C abundance between 2019 and 2020 (p = 0.0004). This aligns with the finding of an increase in schistosome infections in the host snail *Planorbella trivolvis* at the lake (Table 1). In 2019, 46 of 60 (76.7%) sites tested using the species-specific assays (those positive using the pan-avian assay) and 38 of 45 (84.4%) in 2020 tested positive for Avian schistosome C. The mean relative abundance of the positive sites at South Lake Leelanau was in 5.81 +/- 1.00 log_10_ GC/sample in 2019 and in 4.88 +/- 1.29 log_10_ GC/sample 2020. At Walloon Lake, of the sites tested using the species-specific assays, 96 of 119 (80.7%) in 2019 and 60 of 89 (67.4%) in 2020 were positive for this parasite. The mean relative of abundance of Avian schistosome C at positive sites at Walloon Lake was 6.22 +/- 1.42 log_10_ GC/sample in 2019 and 5.87 +/- 1.39 log_10_ GC/sample in 2020. Avian schistosome C displayed two peaks in abundance at both Walloon and South Lake Leelanau, one in early June that persisted through late July, and a second peak in late August, and this trend can be seen in both 2019 and 2020. Similar peaks occur at Glen Lake and North Lake Leelanau; however fewer sites are positive and whole lake abundance of the parasite was lower at these lakes (Figure 2; Figure 3). In 2019, Avian Schistosome C was detected in water samples more frequently at Glen Lake than in 2020, but at a low relative abundance in each sample (Figure 2; Figure S1). Of the sites at Glen Lake that were tested using the species-specific assays, 36 of 119 (30.3%) were positive for Avian schistosome C in 2019 and 17 of 98 (17.4%) in 2020. The mean relative abundance of Avian schistosome C DNA at positive sites in 2019 was 5.14 +/- 1.31 log_10_ GC/sample and 4.65 +/- 1.31 log_10_ GC/sample in 2020. At North Lake Leelanau, of the sites that were tested using the species-specific assays, 14 of 60 (23.3%) in 2019 and 6 of 46 (13.0%) in 2020 were positive for Avian Schistosome C. The mean relative abundance of the sites that were positive for Avian Schistosome C was 4.73 +/- 1.54 log_10_ GC/sample in 2019 and 4.58 +/- 1.55 log_10_ GC/sample in 2020.

**Table 1.**
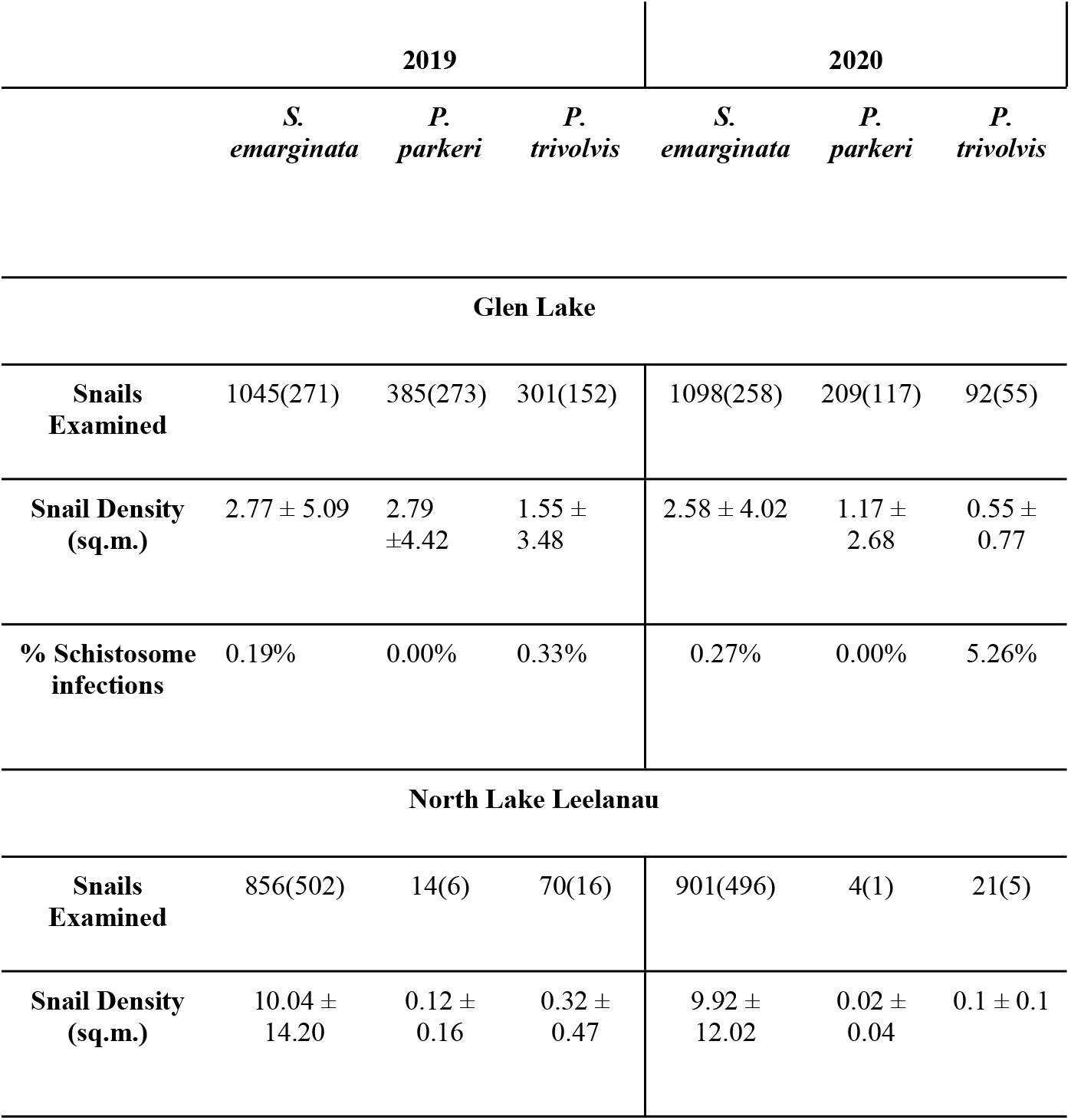
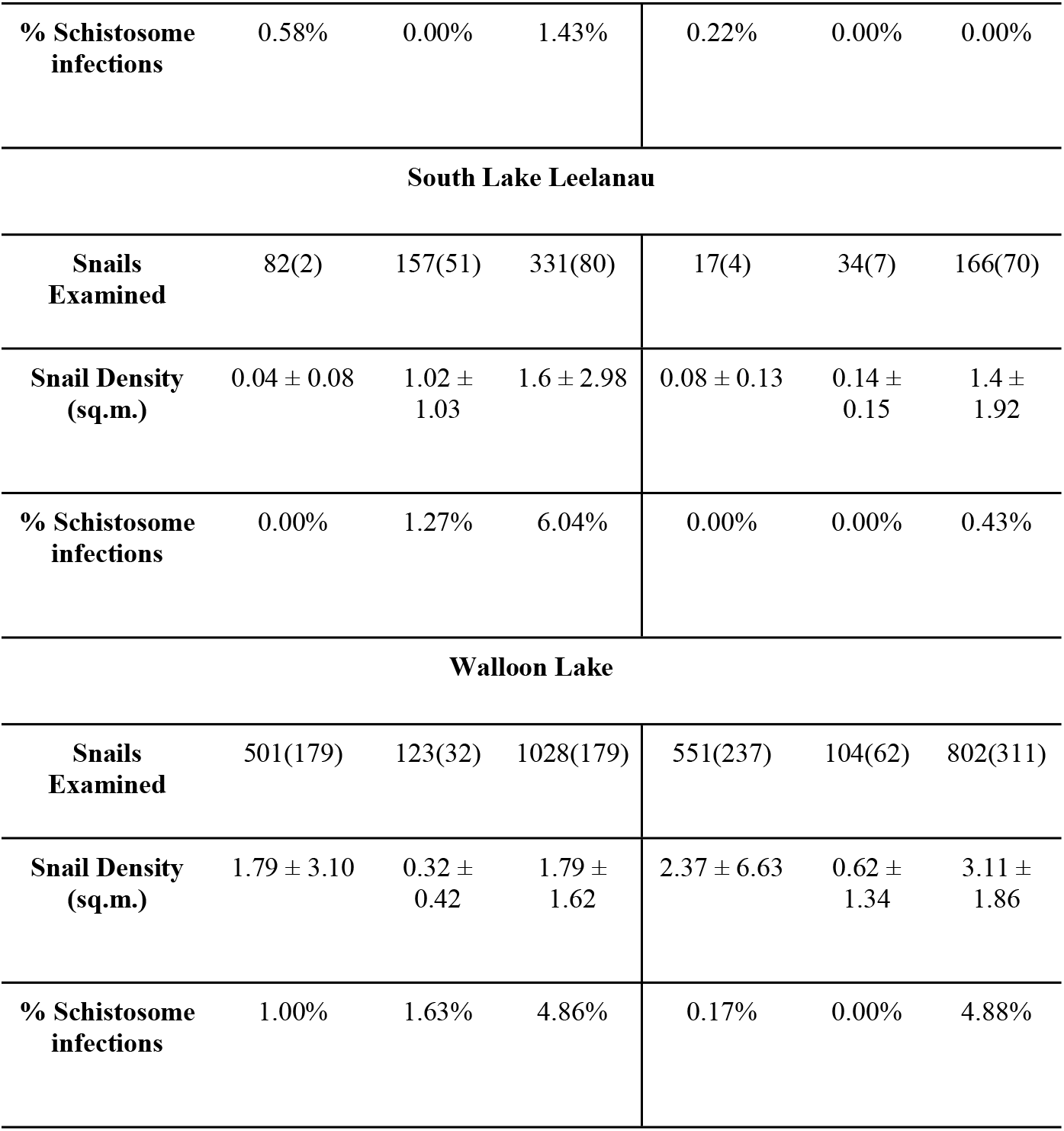
Snail population density (± standard deviation) and avian schistosome infection prevalence for each site at each study lake in 2019 and 2020. Snail density data was collected from 10 m^2^ randomly selected locations at each site during the last two weeks of June. Numbers from the density study are in parentheses beside total snails collected at each site which was used to assess infection prevalence.

### *T. physellae, T. szidati, and A. brantae* are minor contributors to the avian schistosome populations on these lakes

*T. physellae* was detected at all four study lakes, but with low frequency and abundance in 2019 and 2020 (Figure 2, panels I-L; Figure 3). In 2019, every schistosome-positive water sample was also tested for *T. szidati* and *A. brantae* using previously published assays [18]; however, we did not detect these parasites at any of the four lakes in 2019 and did not test for these parasites in 2020.

### Snail density and parasite infection prevalence in snails changed between 2019 and 2020

Snail densities, avian schistosome infection prevalence in snails, and non-avian schistosome infection prevalence in snails were measured in 2019 and again in 2020 at all water sample collection sites (Table S1). North Lake Leelanau had the highest density of *S. emarginata* (intermediate host of *T. stagnicolae*) in both years with a density of 10.04 ± 14.20 snails/m^2^ and 9.92 ± 12.02 snails/m^2^ respectively and had a *T. stagnicolae* infection prevalence of 0.58% in 2019. In 2020 this decreased to 0.22%. Glen lake had the second highest density of *S. emarginata* snails with 2.77 ± 5.09 snails/m^2^ in 2019 and 2.58 ± 4.02 snails/m^2^ in 2020. The *T. stagnicolae* infection prevalence in *S. emarginata* at Glen Lake was 0.19% in 2019, rising to 0.27% in 2020. Both North Lake Leelanau and Glen Lake are lakes that are dominated by *T. stagnicolae* (which utilizes *S. emarginata* as an intermediate host) and both lakes had consistent merganser relocation programs. Walloon Lake had a *S. emarginata* density of 1.79 ± 3.10 snails/m^2^ in 2019, and 2.37 ± 6.63 snails/m^2^ in 2020. The *T. stagnicolae* infection prevalence was 1.00% in 2019, decreasing to 0.17% in 2020. South Lake Leelanau had an *S. emarginata* density of only 0.04 ± 0.08 snails/m^2^, which rose to 0.08 ± 0.13 snails/m^2^ in 2020. The *T. stagnicolae* infection prevalence was 0% in both years. No snails were found to host *T. stagnicolae* at South Lake Leelanau in 2019 or 2020 (Table 1).

### Common merganser relocation does not significantly impact *T. stagnicolae* cercariae abundance

The *T. stagnicolae* DNA abundance data for each lake in 2019 and 2020 (i.e.: total abundance data, all sites and timepoints) was first graphed using box plots, and this visualization showed slight differences between the lakes in both years, but the data had large variation, especially at South and North Lake Leelanau and Walloon Lake. A single factor ANOVA indicated that there were no significant differences between *T. stagnicolae* abundance at each of the four lakes in both 2019 (p = 0.2) and 2020 (p = 0.9). Furthermore, when *T. stagnicolae* DNA abundance was compared between each of the lakes in both years, there were no significant differences found.

The abundance of *T. stagnicolae* was compared between 2019 and 2020 at Glen Lake and North Lake Leelanau, because in 2020 both lakes discontinued their merganser relocation programs to control swimmer’s itch. Whole lake (i.e.: total abundance across all sites and timepoints) was compared in an unpaired two-samples Wilcox test. There was no significant difference between *T. stagnicolae* abundance from 2019 to 2020 at North Lake Leelanau (p = 0.053). However, there was a statistically significant decrease in *T. stagnicolae* abundance at Glen Lake between 2019 and 2020 (p = 0.04). Once again, after hatch-year snails collected in 2020 would reflect trapping and relocation efforts in 2019 since the parasites overwinter in the snail hosts.

### The effect of environmental variables on avian schistosomes

The effect of environmental variables on total cercariae concentrations was assessed using a Hurdle Model. Hurdle models contain two parts, one component (the hurdle part) contains factors that predict zeros (i.e.: no cercariae present in a sample), while the second part of the model predicts non-zero counts. Two predictors were included in the final hurdle-part of the model, temperature and wind speed (Table 2). Lower temperatures predict fewer cercariae in water samples, while high wind speeds also predict fewer cercariae in the water. Every 1 degree increase in temperature increases the odds of having greater than 0 cercariae by 1.60 (p < 0.05), while every 1 km/hr increase in wind speed decreases the odds of having greater than 0 cercariae by 0.88 (p < 0.05). The effects of wind direction, lake, and wind speed, were assessed in the non-hurdle part of the model. Having no wind (i.e., no reading on the handheld anemometer) increased odds of cercariae concentrations by 1.28 times (p<0.05). Every 1 km/hr increase in wind speed predicts a −0.026 (p < 0.01) fold change in cercariae concentrations.

**Table 2.** Assessing the effect of environmental variables on cercariae concentrations throughout the 2019 and 2020 field seasons.

| **Count model (neg bin, log link)** |  |  |  |  |
| --- | --- | --- | --- | --- |
|  | Slope | Odds | Std. Error | Sig. |
| Wind direction relative to the shore |  |  |  |  |
| Reference: Offshore wind |  |  |  |  |
| No Wind | 0.248769 | 1.282445844 | 0.115430 | * |
| Along Shore | 0.125137 | 1.133303924 | 0.112480 |  |
| On Shore | 0.188503 | 1.207441257 | 0.102280 |  |
| Lake |  |  |  |  |
| Reference: South Lake Leelanau (Neg. Control) |  |  |  |  |
| WAL | 0.188247 | 1.207131231 | 0.104171 |  |
| NLL | 0.036631 | 1.037310588 | 0.129946 |  |
| GLEN | 0.114867 | 1.121724780 | 0.108943 |  |
| Wind speed |  |  |  |  |
| Wind speed | -0.025938 | 0.974395295 | 0.009269 | ** |
| Year |  |  |  |  |
| Reference: 2019 |  |  |  |  |
| 2020 | -0.132702 | 0.875725641 | 0.072010 |  |

**Zero hurdle model (censored neg bin, log link)**
|  | Slope | Odds | Std. Error | Sig. |
| --- | --- | --- | --- | --- |
| Temperature | 0.47280 | 1.604480782 | 0.19528 | * |
| Wind speed | -0.13133 | 0.876930631 | 0.06262 | * |

Significance: \* 0.05 \*\* 0.01 \*\*\* 0.0001
Theta: count= 3645096.6118, zero = 0.15 on 14 degrees of freedom.

## Discussion

The avian schistosome life cycle was first described in Michigan in 1928 [20]. Since this discovery, the state has implemented various strategies to control swimmer’s itch over the past 80 years. In the 1980s, studies were conducted to gauge the efficacy of treating mallards (*Anas platyrhynchos*) and common mergansers (*M. merganser*) with Praziquantel, an anthelmintic drug that is also used to treat human schistosomiasis. These studies suggested Praziquantel was effective at reducing the infection prevalence of schistosomes in birds and indicated that yearly treatment of birds at specific sites may reduce swimmer’s itch in localized areas [8,9]. These studies led to the waterfowl-based control efforts that are outlined in this study, that focus on lake-wide relocation of a top-tier predatory bird from numerous inland Michigan lakes. While the principle of parasite life cycle interruption by host relocation seems sound in theory, here we present evidence that suggests that lake-wide relocation of resident common mergansers had little to no measurable impact on the population of *T. stagnicolae* on any of the inland lakes that participated in our comprehensive study.

### A resident definitive host population is not required to maintain a large avian schistosome population

We found no statistically significant differences between *T. stagnicolae* abundance between lakes with resident common merganser control programs (Glen and North Lake Leelanau), the negative control lake (Walloon Lake) which had resident common mergansers present on the lake in 2018-2020, or the positive control lake (South Lake Leelanau) which does not support resident mergansers. There was not a significant difference between Glen Lake and Walloon Lake in 2019, nor in 2020. If merganser relocation were effective, we would have expected the cercariae abundance at Glen Lake to be similar to that of South Lake Leelanau, and to differ from Walloon Lake.

In 2020, Glen and North Lake Leelanau discontinued relocating summer resident common merganser broods. We compared *T. stagnicolae* cercariae abundance in 2019 and 2020 and found that Glen Lake had a moderately statistically significant reduction in *T. stagnicolae* abundance when summer resident common mergansers remained onto the lake. However, since schistosomes over-winter in snails and we only collected after hatch-year snails, cercariae data reflect previous year infection logistics.

Furthermore, lakes with and without control programs show very similar longitudinal abundance of *T. stagnicolae* throughout the year. *T. stagnicolae* concentrations across the whole lake rise sharply in early June (Figure 2; Figure 3). The results of the hurdle model (Table 2), suggest this effect is likely due to water temperatures at this time of the year becoming conducive to cercariae emergence from snails. This observation corroborates findings from previous laboratory studies that have found cercariae emergence to be temperature dependent in numerous digenean trematode systems [21–24]. *T. stagnicolae* abundance remains high until late September at lakes with and without control programs (Figure 3).

This study shows that *T. stagnicolae* maintains its life cycle in the absence of a continuous definitive host presence for most of the transmission season. Our group collected additional field data in summer 2020 on Higgins Lake (Roscommon Co., MI) that would support this study. Not only were all merganser broods removed since 2015, but they also systematically eliminated adult and breeding mergansers through two years of spring merganser duck hunts (2015 and 2016), filling nesting holes, and by euthanizing breeding hens for ancillary research projects. This resulted in a stable summer resident merganser population of 9 broods in 2015 to 0 broods in 2020. Even with these definitive host mitigation efforts, numerous cases of swimmer’s itch were reported in 2020 and average levels of *T. stagnicolae* were found in the water in July [25].

If we assume that the common merganser is the only suitable definitive host for *T. stagnicolae* at inland Northern Michigan lakes, as is suggested by controlled challenge studies [7], then migratory common mergansers in the spring and fall must have been contributing enough parasites into the ecosystems of Glen Lake and South and North Lake Leelanau to sustain *T. stagnicolae* populations in the intermediate host *S. emarginata* until fall, when the migratory birds return. Since all broods were trapped <4 weeks from hatching, ducklings could not have contributed to the parasite load on lakes implementing merganser control. In addition, non-nesting adult mergansers usually vacate these inland lakes by early-mid summer, thereby reducing the likelihood of them contributing to parasite transmission. Thus, birds that are not resident contributors to the biodiversity of a specific lake ecosystem nonetheless can contribute and sustain the parasite biodiversity of an ecosystem in the case of this avian schistosome species. Underpinning these observations are our observations of the intermediate snail host community. North Lake Leelanau, Glen Lake and Walloon Lake have higher densities of *S. emarginata* than South Lake Leelanau. It appears that lakes with relocation programs (i.e., Glen Lake and North Lake Leelanau) that have higher snail densities will still have high cercariae concentrations, due to the exponential amplification of the parasite within the intermediate host. Similarly, this exponential amplification explains why South Lake Leelanau, which has a low *S. emarginata* density and does not have resident birds, still has *T. stagnicolae* cercariae in the water. We also noted decreases in *S. emarginata* density at Glen Lake and North Lake Leelanau between 2019 and 2020, whereas the population of this snail increased at South Lake Leelanau and Walloon Lake. However, there was not a corresponding increase in snail infection prevalence. Changes in the snail populations undoubtably underpin changes in cercariae abundance from year to year, and future research from our group will explore how changes in *S. emarginata* snail density at these lakes over the course of three years are related to changes in *T. stagnicolae* parasite populations.

### Avian schistosome C is an important contributor to avian schistosome populations in northern Michigan Lakes, while *T. physellae* is a minor contributor

In 2018 our group discovered a novel avian schistosome-like species emerging from a *P. trivolvis* (*=Helisoma trivolvis*) snail. As reported in McPhail *et al*. [19], this proved to be a novel genus of avian schistosome which utilizes definitive host *Branta canadensis* (and possibly others) and caused dermatitis under laboratory conditions. Avian Schistosome C made up a significant component of the avian schistosome community and was the most abundant avian schistosome at Walloon Lake and South Lake Leelanau (Figure 2; Figure 3). This parasite is detectable earlier and persists later into the season than *T. stagnicolae*. Interestingly, lakes with control programs aimed at controlling *T. stagnicolae* have lower abundance of Avian Schistosome C. These lakes also have lower densities of *P. trivolvis*, the intermediate host of Avian Schistosome C. As might be expected, South Lake Leelanau and Walloon Lake have higher densities of *P. trivolvis* and consequently higher relative abundances of Avian Schistosome C.

*Trichobilharzia physellae* was sporadically detected throughout the season at all four lakes. The intermediate host of *T. physellae* in Northern Michigan lakes is *Physellae parkeri*, and the definitive hosts are mallard ducks and common mergansers. Lakes in our study that trapped and relocated summer resident common mergansers have low abundance and more sporadically appearing *T. physellae* populations than the control lakes. However, the control lakes also support low abundances of this parasite. Interestingly, Glen Lake has a high density of *P. parkeri*, despite having a low reactive abundance of *T. physellae*. Because we do not know the prevalence of this parasite in bird populations, it is possible this parasite has a lower infection prevalence in birds and consequently could produce lower percent abundance in water samples.

### Wind, Waves, Temperature and Location Affect Cercariae Concentrations

This study and others have assessed the effect of environmental conditions on cercariae concentrations and incidence of swimmer’s itch [14,16]. Sckrabulis *et* al. [14] indicates that onshore winds represent an increased risk for higher cercariae concentrations, and that onshore winds increase the daily incidence of swimmer’s itch. While we found that onshore winds do increase the odds of cercariae concentrations by 1.21 times, this was not statistically significant. This can likely be explained by how the water samples were collected for this study. By sampling along the length of a dock, rather than solely along the shore, it is likely that the effect of onshore winds on cercariae concentration was diluted.

This study also shows that temperature and wind speed act as “hurdles” predicting an absence of cercariae at beaches (Table 2). Numerous laboratory studies have demonstrated that cercariae emergence is temperature dependent [22,26,27], and as such it is unsurprising that temperatures less than 15°C are predictive of 0 cercariae detected in the water. However, temperature was not a significant predictor of non-zero values in our model. While temperature has been extensively demonstrated to drive cercariae survival and production [27], this effect varies between different species of trematode and their snail hosts. Therefore, our pan-avian qPCR approach to measuring cercariae in the water would mask this effect.

The results of temperature and their effect on cercariae concentrations is corroborated in Sckrabulis *et al* [14]. These authors did not find temperature to be predictive of increases in swimmer’s itch incidence. Windspeed was a significant predictor in our model, with increased wind speed resulting in a decrease in cercariae concentrations at local beaches. In addition, strong winds likely congregate the floating cercariae very near shore (<1m) which was inside our sampling zone. This study also highlights the variable nature of cercariae concentrations at individual locations (Figure 1; Figure 2), and highlights that the risk of encountering cercariae in water samples differs spatially and temporally at locations across a lake, with some sites having relatively low concentrations of cercariae, while others are nearly constantly inundated with cercariae.

### Implications for Swimmer’s itch Management

The results of this study demonstrate definitively that the relocation of *M. merganser* from lakes in northern Michigan has not been effective at reducing or controlling swimmer’s itch causing parasite *T. stagnicolae* at our study lakes. These results are not surprising and have been demonstrated in other analogous host parasite systems, notably that of human schistosomiasis. Sokolow *et al*. [4], assessed the effectiveness of different control programs aimed at eliminating *S. mansoni* in human populations and found that intermediate host control (i.e., snail control) was most effective at eliminating *S. mansoni* from communities, while mass drug administration, a control effort aimed at eliminating the parasite from the definitive human host, was less effective. Taken together, what these results suggest is that definitive host control is not alone sufficient to interrupt the life cycle of a schistosome (whether avian or mammalian) and thereby stop parasite transmission in an area unless it is absolute. Our results show that transient introductions of avian schistosomes from migratory common mergansers is sufficient to sustain infections in the snail intermediate hosts throughout the summer, even when resident mergansers have been removed from lakes for 3 consecutive years. Birds migrating along flyways are likely making meaningful contributions to the parasite communities at their landing sites. This should be considered along with the likely substantial impact that relocation of a top predator has in these inland lake ecosystems, when considering future merganser relocation programs to control for swimmer’s itch. Moreover, while there have been notable exceptions [28–30], mathematical models and the consensus of scientific literature suggest that high host diversity typically decreases disease risk [31–35]. Future research in Michigan will focus on understanding whether this principle holds true for swimmer’s itch-causing parasites, measuring whether changes in bird and snail diversity lead to measurable impacts on parasite diversity.

This study also provides an interesting observation for future ecological studies. Previous work has shown that definitive bird hosts drive the composition of the trematode community, and trematodes have been utilized in wetland restoration studies as surrogates for bird host diversity [36–38]. This study suggests that migrant birds make an important and sustained contribution to the trematode community. Trematodes could provide an imprint of transient host presence and could be an interesting sentinel measurement of true biodiversity for ecosystems. Surveys of trematode diversity may also provide information on transitory host presence in a way that traditional bird survey methods may not capture.

In conclusion, the swimmer’s itch problem in Michigan is complex, and has resulted in researchers and riparians alike searching for a solution for the better part of the last century. Even with the long history of research in this area, there continue to be discoveries. These include the presence of a novel avian schistosome species found in the state that has been revealed as another major contributor to the schistosome community at these lakes. Furthermore, with this study, we demonstrate that removing the resident definitive host of the dominant avian schistosome, *T. stagnicolae*, from the lakes did not have the hypothesized effect of interrupting the life cycle and reducing the numbers of this parasite found in the water. As such, swimmer’s itch research is ongoing, and future research will continue to focus on these avian schistosomes, and the contributions made to the schistosome community by non-resident migrants. In addition, we predict a practical paradigm shift away from whole-lake control to individual prevention will occur, and we plan to assess the myriad prevention strategies already employed by riparians. This new paradigm will empower individual riparians, reduce costs for lake associations, and have little negative ecological impact.

## Methods

### Study Locations and Sampling

Sampling sites were selected in coordination with community-based partners. The sites were selected to give broad coverage around the lake, to target high use swimming areas or areas with perennial swimmer’s itch problems, and for easy access to ensure safety and convenience for volunteers. Ten (10) sites were chosen on both Glen and Walloon Lake, and 5 sites each on North and South Lake Leelanau (Figure 4).

**Figure 4:**
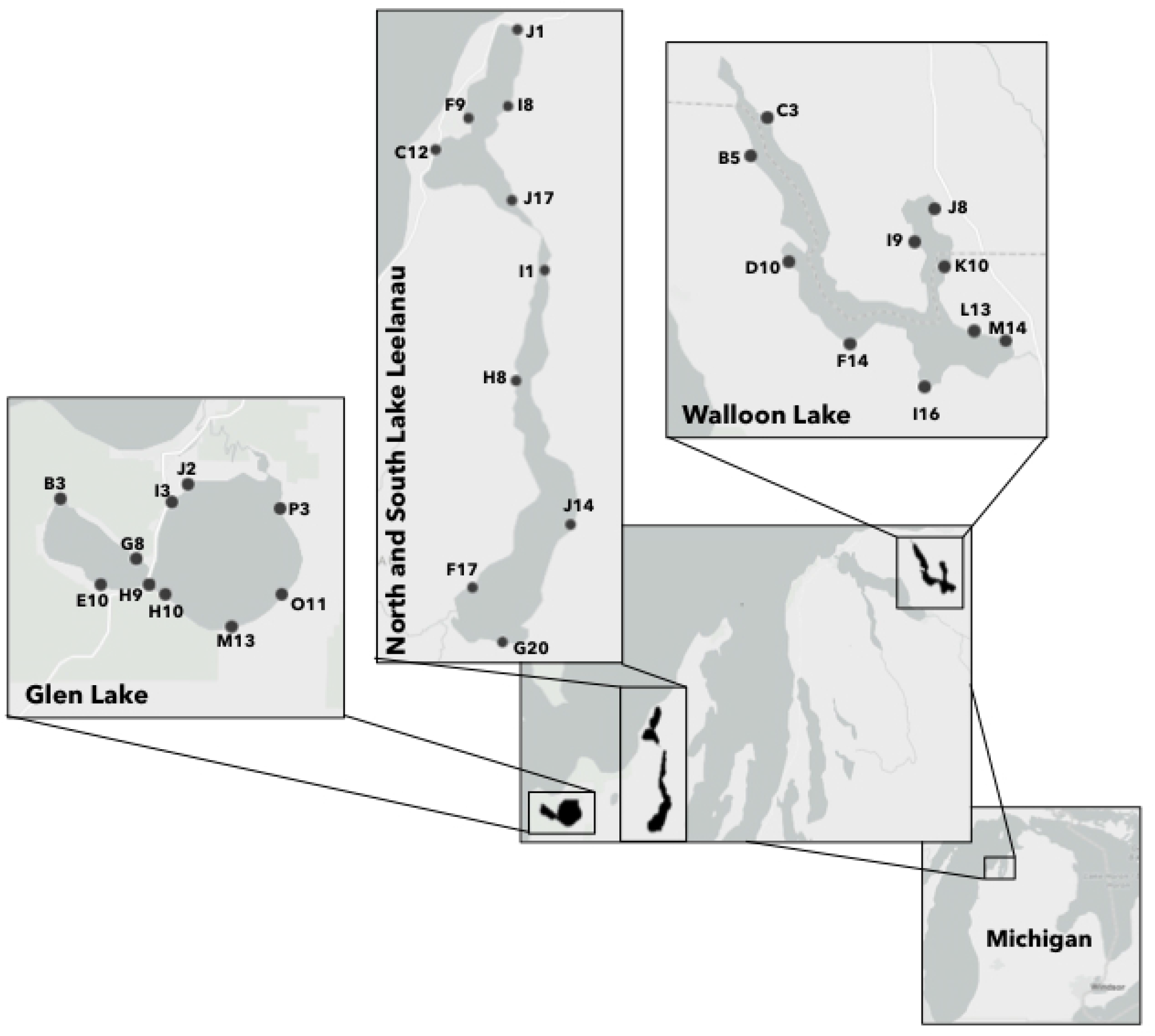
Sampling locations in on Glen Lake, North and South Lake Leelanau, and Walloon Lake.

Community partners collected samples every Tuesday morning from May 21 through October 22 between 8 a.m. and noon. Each team of samplers was supplied with a standardized sample collection kit that contained: a 20 μm mesh plankton tow, floating thermometer, handheld anemometer, 1-liter scoop, sterile 50 mL sampling tubes, 95% ethanol, and a laminated instruction sheet. All volunteers were given onsite training to ensure proper water collection protocol was followed. In addition, an instructional video was created for their review. Sampling was conducted by passing 25 L of water through the 20 μm plankton tow. Water was collected one litre at a time along a swath of beach with average depth of approximately 0.5 m by sampling along the length of a dock or wading into the water. This method of water collection has been previously described for use in performing qPCR cercariometry [16–18].

Water samples were reduced to approximately 25 mL and then preserved by adding approximately 25 mL of 95% ethanol. Volunteers transported samples the same day to the central laboratory. Samples were then refrigerated until processing (usually within 5 days). The 50 mL samples were suction filtered through a 0.4 μm filter (Pall FMFNL1050) from which DNA was extracted.

Meta-data were collected at each site on each collection day. Average wind speed was recorded using a handheld anemometer. Wind direction and relative water accumulation (onshore, offshore, alongshore) were recorded as observations. Water temperature was recorded to the nearest 0.1 degree C using a floating pool thermometer. Date, location, time of water collection, and general weather conditions (clear, partly cloudy, etc.) were also recorded.

### DNA extraction and qPCR

DNA extraction was accomplished using the Qiagen (USA) DNEasy Blood and Tissues kit as described in [16]. In short, the tissue lysis protocol was followed but with one modification; filter membranes were bead-beat using a vortex adapter for 5 minutes with approximately 100 μL of 0.1mm diameter glass beads (BioSpec #11079101) prior to chemical lysis [18]. The qPCR cercariometry for all assays were carried out using the following thermocycling parameter: QuantStudio 3 (Thermo Fisher Scientific, USA) using a standard, 40 cycle, two-step reaction. The thermocycling parameters were a 30 second hold at 95°C, followed by a 30 second denaturation cycle at 95°C, and a 60°C annealing cycle. Each qPCR reaction had a final volume of 20μL, and we added 5μL of DNA to each reaction. Inhibition controls were performed on samples according to the procedure outlined in previous qPCR cercariometry papers [16,18].

### Primers and Probes

qPCR assays for the pan avian schistosome qPCR assay, and the species-specific *T. stagnicolae*, and *T. physellae* assays were previously published [15,16,18]. The sequences for these assays are listed in Table S1.

### Development of a species-specific qPCR Avian Schistosome C

A cytochrome *c* oxidase subunit 1 (CO1) gene-targeting assay specific for Avian Schistosome C was developed. Multiple specimens visually identified as Avian Schistosome C shed from snails were collected from Michigan in 2018 and 2019 and underwent barcoding of the Folmer region [39,19]. The sequences generated were aligned to find a conserved region of about 100 nucleotides in length that would be an appropriate diagnostic target. The target was also aligned against related avian schistosomes to ensure there would not be cross reactivity (Figure S2). Avian Schistosome C CO1 sequence most closely resembled *T. stagnicolae, T. physellae, T. franki and T. querquedulae*, and therefore the alignment focuses on these species. Cross reactivity was also tested experimentally against *T. physellae, T. stagnicolae, T. szidati* and *Anserobilharzia brantae* and no cross reactivity was observed. The LOD95 of this assay is 89.9 copies/5uL (upper limit (with 95% confidence): 181 copies/5uL, lower limit (with 95% confidence): 44.6 copies/5uL). The primer and probe sequences of this assay are listed in Table S1.

### Longitudinal assessment of three species of avian schistosomes in four inland lakes in northern Michigan

We utilized three avian schistosome source tracking assays (*T. stagnicolae, T. physellae* and Avian Schistosome C) to conduct a longitudinal analysis of the avian schistosome community at four lakes in 2019 and 2020. Water samples were first screened via the *18s rDNA* pan avian schistosome assay (Figure S2), and samples negative via this assay were excluded from being tested using the species-specific assays. Walloon Lake, South Lake Leelanau, North Lake Leelanau, and Glen Lake were assessed. Walloon and South Lake Leelanau served as control lakes. Walloon lake does not, and has never had, a merganser relocation control program, and has numerous resident merganser broods on the lake each summer. South Lake Leelanau also does not, and has never had, a merganser removal control program, but this lake does not support summer resident common mergansers as evidenced by our whole-lake waterfowl surveys which have been conducted annually since 2016. Residents still report swimmer’s itch at this lake. Glen Lake and North Lake Leelanau (which is separated from South Lake Leelanau by a riverine system called The Narrows), both had merganser removal control programs each year beginning in 2017. A total of 455 individual mergansers were trapped and relocated from these two lakes from 2017-2019. In 2020, merganser relocation programs ended at Glen Lake and North Lake Leelanau.

### Snail collections and density assessments

Snails were collected between 19 June and 29 June, 2019 and 2020 at the 30 sampling sites on all four lakes. To assess relative abundance, 1m^2^ weighted hoops were randomly tossed throughout the collection site. Divers in full wetsuits and snorkeling gear collected all snails of all species within the hoops using specially designed snail scoops with attached mesh bag nets.

The collected snails were placed in labeled buckets filled with fresh lake water and transported to the lab. Snails were sorted morphologically, counted, and each snail placed in its own compartment of a 12-well culture plate partially filled with conditioned water. The snails were exposed to natural lighting until dark, kept in the dark until dawn, and then exposed to natural and artificial light for at least an hour before each snail was examined for shedding cercariae using a dissection microscope (10x). For those snails that shed cercariae, the cercariae were morphologically characterized and those that were avian schistosomes were counted.

### Statistics

Heat maps and bar charts were created using GraphPad Prism version 9.2 for Mac OS, GraphPad Software, San Diego, California, USA, and maps were created using ArcMap (Esri, USA). A hurdle count regression model using a negative binomial distribution and a log link function was used to assess the effect of various environmental variables on cercariae concentrations. Hurdle models are two-part models that specify one process for zero counts, and one process for nonzero counts (i.e.: positive samples arise from crossing a hurdle). This approach recognizes that theoretically different processes may contribute more to the absence of cercariae than their presence. The hurdle model was therefore selected both on a theoretical basis, but also because the data were over-dispersed, and zero inflated. The poisson, quasi-poisson, negative binomial, and zero inflated poisson models were also tested, but did not fit the data as well as the Hurdle model based on AIC criterion (where possible). Additionally, the Hurdle model resulted in lower dispersion parameters, smaller standard deviations and an improved log likelihood. These models were implemented in R 4.0.0 [40] using the pscl, MASS, and psych packages [41–43].

## Author contributions

**SPR**: Data Curation, Formal Analysis, Investigation, Methodology, Project Administration, Resources, Software, Validation, Writing – Original Draft. **BAM**: Data Curation, Formal Analysis, Investigation, Resources, Software, Writing – Review & Editing. **RLR**: Conceptualization, Data Curation, Project Administration, Resources, Methodology, Writing – Review & Editing. **KF**: Data Curation, Resources, Writing – Review & Editing. **AT**: Investigation, Software. **PCH**: Conceptualization, Funding Acquisition, Methodology, Project Administration, Supervision, Visualization, Writing – Editing & Review.

## Acknowledgements

This work was not possible without the dedicated efforts of many riparian water collectors who collected samples every Tuesday morning from May to October for two years. Volunteers from Glen Lake: Joe Blondia, Dale DeJager, Andy DuPont, Cecelia Denton, Evan Fink, Ed Gergosian, Rob Karner, Bill Meserve, Shelley Walter, Holly Wright. Volunteers from South Lake Leelanau: Dan Harkness, Thad Popa. Volunteers from North Lake Leelanau: Jeff Green, Brian Price, Jim Wysor. Volunteers from Walloon Lake: Connor Dennis, Betony Braddock, Jac Talcott, John Marklewitz, Russel Kittleson, Mary Pat Goldich. We also acknowledge the many hours of work spent collecting snails at all 30 locations by Chris Froelich, Daniel Clyde, and Matt Schuiling. This work was supported by funding from the Natural Sciences and Engineering Council of Canada (NSERC) # <u>2018-05209</u> and <u>2018-522661</u> (PCH), and by Alberta Innovates <u>#2615</u> (PCH).

---

**Table S1:**
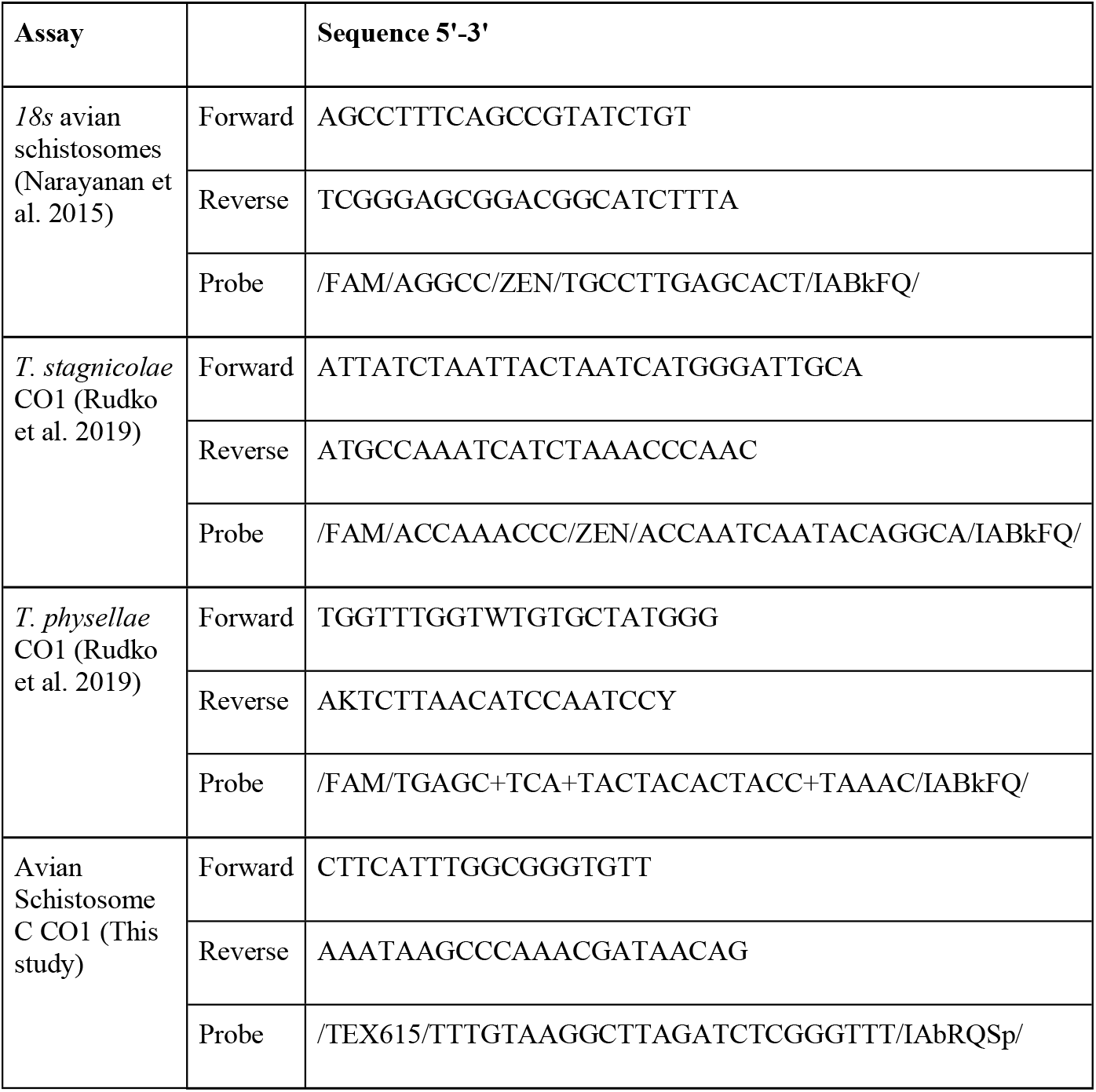
Primers and Probes

Figure S1: Accumulated Log_10_ transformed cercariae/25L combining 2019 and 2020 data for each lake (A: Glen Lake; B: Walloon Lake; C: North Lake Leelanau; D: South Lake Leelanau). Combined 2019 and 2020 proportions for each avian schistosome species (red = Avian schistosome sp. C, blue = *T. stagnicolae*, green = *T. physellae*) is presented in the lower bar charts.

Figure S2: Alignment of the qPCR region of CO1 that is targeted by the species-specific qPCR tests.

## References

1. World Health Organization. Weekly Epidemiological Record. 2019 p. 94, 601–12. Report No.: 50.

2. Soldánová M, Selbach C, Kalbe M, Kostadinova A, Sures B. Swimmer’s itch: etiology, impact, and risk factors in Europe. Trends Parasitol. 2013;29(2):65–74.

3. Horák P, Mikeš L, Lichtenbergová L, Skála V, Soldánová M, Brant SV. Avian schistosomes and outbreaks of cercarial dermatitis. Clin Microbiol Rev. 2015;28(1):165–90.

4. Sokolow SH, Wood CL, Jones IJ, Swartz SJ, Lopez M, Hsieh MH, et al. Sokolow SH, Wood CL, Jones IJ, Swartz SJ, Lopez M, Hsieh MH, et al. (2016) Global assessment of schistosomiasis control over the past century shows targeting the snail intermediate host works best. PLoS Negl Trop Dis. 2016;10(7):e0004794.

5. Wood CL, Summerside M, Johnson PTJ. How host diversity and abundance affect parasite infections: Results from a whole-ecosystem manipulation of bird activity. Biol Conserv. 2020;248:108683.

6. Michigan Department of Natural Resources Wildlife Division. Common Merganser Control Policy and Procedures [Internet]. Michigan; 2018 Feb p. 7. (Wildlife Policy and Management). Report No.: 08.01.17. Available from: https://www.watershedcouncil.org/uploads/7/2/5/1/7251350/mdnr_come_policy_signed_02282018.pdf

7. Brant SV, Loker ES. Molecular systematics of the avian schistosome genus Trichobilharzia (Trematoda: Schistosomatidae) in North America. J Parasitol. 2009;95(4):941–63.

8. Reimink RL, DeGoede JA, Blankespoor HD. Efficacy of Praziquantel in natural populations of mallards infected with avian schistosomes. J Parasitol. 1995;81(6):1027–9.

9. Blankespoor CL, Reimink RL, Blankespoor HD. Efficacy of Praziquantel in treating natural schistosome infections in common mergansers. J Parasitol. 2001;87:424–6.

10. Pardieck KL, Ziolkowski Jr. DJ, Lutmerding M, Hudson M-AR. North American breeding bird survey dataset 1966 - 2017 version 2017.0., [Internet]. U.S. Geological Survey, Patuxent Wildlife Research Center; 2018. Available from: https://doi.org/10.5066/F76972V8.

11. State of Michigan Department of Natural Resources. Common Merganser Control Permit and Regulations. 2018.

12. Hoeffler DF. Cercarial dermatitis: its etiology, epidemiology, and clinical aspects. Arch Environ Health. 1974;29(4):225–9.

13. Selbach C, Soldánová M, Sures B. Estimating the risk of swimmer’s itch in surface waters - A case study from Lake Baldeney, River Ruhr. Int J Hyg Environ Health. 2016;219:693–9.

14. Sckrabulis JP, Flory AR, Raffel TR. Direct onshore wind predicts daily swimmer’s itch (avian schistosome) incidence at a Michigan beach. Parasitology. 2020;147:431–40.

15. Narayanan J, Mull BJ, Brant SV, Loker ES, Collinson J, Secor WE, et al. Real-time PCR and sequencing assays for rapid detection and identification of avian schistosomes in environmental samples. Appl Environ Microbiol. 2015;81(12):4207–15.

16. Rudko SP, Reimink RL, Froelich K, Gordy MA, Blankespoor CL, Hanington PC. Use of qPCR-based cercariometry to assess swimmer’s itch in recreational lakes. EcoHealth. 2018;15(4):827–39.

17. Froelich K, Reimink RL, Rudko SP, VanKempen AP, Hanington PC. Evaluation of targeted copper sulfate (CuSO4) application for controlling swimmer’s itch at a freshwater recreation site in Michigan. Parasitol Res. 2019;118(5):1673–7.

18. Rudko SP, Turnbull A, Reimink RL, Froelich K, Hanington PC. Species-specific qPCR assays allow for high-resolution population assessment of four species avian schistosome that cause swimmer’s itch in recreational lakes. Int J Parasitol Parasites Wildl. 2019;

19. McPhail BA, Rudko SP, Turnbull A, Gordy MA, Reimink RL, Clyde D, et al. Evidence of a putative novel species of avian schistosome infecting Planorbella trivolvis. J Parasitol. 2021;107(1):89–97.

20. Cort WW. Schistosome dermatitis in the United States (Michigan). J Am Med Assoc. 1928;90:1027–9.

21. Shiff CJ, Evans A, Yiannakis C, Eardley M. Seasonal influence on the production of Schistosoma haematobium and S. mansoni cercariae in Rhodesia. Int J Parasitol. 1975;5(1):119–23.

22. Lo C-T, Lee K-M. Pattern of emergence and the effects of temperature and light on the emergence and survival of heterophyid cercariae (Centrocestus formosanus and Haplorchis pumilio). J Parasitol. 1996;82:347–50.

23. Abrous M, Rondelaud D, Dreyfuss G. Influence of low temperatures on the cercarial shedding of Paramphistomum daubneyi from the snail Lymnaea trunculata. Parasite. 1999;6(85–88).

24. Haas W, Beran B, Loy C. Selection of the host’s habitat by cercariae: from laboratory experiments to the field. J Parasitol. 2008;94:1233–8.

25. Higgins Lake Foundation Final Report. 2020. Available from: https://secureservercdn.net/104.238.71.250/9vm.8cc.myftpupload.com/wp-content/uploads/2020/11/2020-Swimmers-Itch-assessment.pdf

26. Haas W. Physiological analyses of host-finding behaviour in trematode cercariae: adaptations for transmission success. Parasitology. 1994;109:S15–29.

27. Poulin R. Global warming and temperature-mediated increases in cercarial emergence in trematode parasites. Parasitology. 2006;132(01):143–51.

28. Norman R, Bowers RG, Begon M, Hudson PJ. Persistence of tick-borne virus in the presence of multiple host species: Tick reservoirs and parasite mediated competition. J Theor Biol. 1999;200(1):111–8.

29. Gilbert L, Norman R, Laurenson KM, Reid HW, Hudson PJ. Disease persistence and apparent competition in a three-host community: An empirical and analytical study of large-scale, wild populations. J Anim Ecol. 2001;70(6):1053–61.

30. Dobson A. Population dynamics of pathogens with multiple host species. In: American Naturalist. S5 ed. Chicago: The University of Chicago; 2004.

31. Van Buskirk J, Ostfeld RS. Controlling Lyme disease by modifying the density and species composition of tick hosts. Ecol Appl. 1995;5(4):1133–40.

32. LoGiudice K, Ostfeld RS, Schmidt KA, Keesing F. The ecology of infectious disease: Effects of host diversity and community composition on Lyme disease risk. Proc Natl Acad Sci U S A. 2003;100(2):567–71.

33. Ruedas LA, Salazar-Bravo J, Tinnin DS, Armién B, Cácaeres L, García A, et al. Community ecology of small mammal populations in Panamá following an outbreak of Hantavirus pulmonary syndrome. J Vector Ecol. 2004;29(1):177–91.

34. Keesing F, Holt RD, Ostfeld RS. Effects of species diversity on disease risk. Ecol Lett. 2006;9:485–98.

35. Johnson PTJ, Preston DL, Hoverman JT, LaFonte BE. Host and parasite diversity jointly control disease risk in complex communities. Proc Natl Acad Sci. 2013;110:16916–21.

36. Huspeni TC, Lafferty KD. Using larval trematodes that parasitize snails to evaluate a saltmarsh restoration project. Ecol Appl. 2004;14(3):795–804.

37. Hechinger RF, Lafferty KD. Host diversity begets parasite diversity: bird final hosts and trematodes in snail intermediate hosts. Proc R Soc B Biol Sci. 2005;272(1567):1059–66.

38. Huspeni TC, Hechinger RF, Lafferty KD. Trematode Parasites as Estuarine Indicators: Opportunities, Applications, and Comparisons with Conventional Community Approaches. In: Estuarine Indicators. Boca Raton: CRC Press; 2005. p. 297–314.

39. Folmer O, Black M, Hoeh W, Lutz R, Vrijenhoek R. DNA primers for amplification of mitochondrial cytochrome c oxidase subunit I from diverse metazoan invertebrates. Mol Mar Biol Biotechnol. 1994;3(5):294–9.

40. R Development Core Team. R: A language and environment for statistical computing. 2020.

41. Zeilis A, Kleiber C, Jackman S. Regression Models for Count Data in R. 2008.

42. Jackman S. pscl: classes and methods for R developed in the political science computational laboratory [Internet]. United States Studies Centre, University of Sydney, Sydney, New South Wales, Australia; 2017. Available from: https://github.com/atahk/pscl/

43. Revelle W. psych: Procedures for psychological, psychometric, and personality research. In: Compr. R Arch. Netw. (CRAN).; 2018.

